# A glycerol oxidase from Norway spruce (*Picea abies* L. Karst.) expands biochemical and structural attributes of the GMC oxidoreductase superfamily

**DOI:** 10.1101/2025.01.06.631446

**Authors:** Tanja Paasela, Naike Schwenner, Hongbo Zhao, Mengyi Sun, Topi Haataja, Johanna Karppi, Aleksia Vaattovaara, Emma Master, Mats Sandgren, Anna Kärkönen, Maija Tenkanen

## Abstract

- Investigations to uncover the versatility of carbohydrate active enzymes belonging to the auxiliary activity family 3 (AA3) have focused on microbial enzymes. Plant genomes also harbor AA3 encoding genes, and reverse genetics approaches have shown their importance in plant development. To date, however, detailed biochemical characterizations and structures of plant AA3 enzymes have not been reported.
- Here, we describe screening AA3 encoding genes from Norway spruce and the first in-depth biochemical and structural characterization of a plant AA3 oxidase, *Pa*AOX1. *Pa*AOX1 was crystallized, and the structure was solved by X-ray crystallography.
- *Pa*AOX1 demonstrated highest oxidase activity against glycerol. It oxidized glycerol to glyceraldehyde, then further to glyceric acid, and showed preference for L-glyceraldehyde over D-glyceraldehyde. Despite the low sequence similarity with fungal alcohol oxidases, it had all the structural features of the typical AA3 enzyme.
- PaAOX1 has a novel activity among the characterized AA3s, and it showed a new structural arrangement of the region surrounding the catalytic center. Our findings contribute to a deeper understanding of the structural and functional aspects of enzymes in the GMC superfamily, and especially of alcohol oxidases, expanding the knowledge from fungi to plants.

## Introduction

Glucose–methanol–choline (GMC) oxidoreductase superfamily enzymes are flavin adenine dinucleotide (FAD) dependent enzymes that belong to the AA3 family, according to the Carbohydrate– Active enZYmes (CAZy) database (Drula et al., 2022). Some members of this versatile enzyme family have been both structurally and biochemically characterized, mainly from fungal species. GMC superfamily enzymes are also present in bacteria, insects, and plants, albeit much less studied in these taxons (Iida et al., 2007; Kostelac et al., 2022; Kurdyukov et al., 2006). One of the key biological functions of AA3 enzymes in fungi is their involvement in degradation of lignocellulosic plant biomass by improving the activity of other CAZymes, such as peroxidases, laccases, and lytic polysaccharide monooxygenases (LPMOs), by providing them hydrogen peroxide and hydroquinones (Bissaro et al., 2018; Higasi & Polikarpov, 2023; Manavalan et al., 2021; Østby et al., 2022).

The fungal AA3 family enzymes are divided into four subfamilies based on their substrate specificities. Subfamily AA3_1 contains enzymes with cellobiose dehydrogenase activities; subfamily AA3_2 includes aryl-alcohol and glucose oxidases/dehydrogenases, and pyranose dehydrogenases; AA3_3 comprises alcohol oxidases, and subfamily AA3_4 includes pyranose 2-oxidases. Enzymes that prefer molecular oxygen as the electron acceptor are defined as oxidases and those that prefer other electron acceptors, such as metal ions, quinones or phenoxy radicals, are defined as dehydrogenases. The catalytic reaction mechanism of these enzymes can be divided into a reductive phase and an oxidative phase (Hallberg et al., 2003; Sützl et al., 2019; Wongnate & Chaiyen, 2013). In the reductive phase, the substrate is oxidized via a hydride transfer or a radical mechanism, resulting in a reduced FAD and a protonated catalytic histidine. In the second reaction phase, the electrons are transferred to the electron acceptor. Concomitantly, the FAD-containing enzyme is deprotonized to its oxidized state.

Although genes encoding putative AA3 family enzymes are present in plant genomes, only a few published reports about these enzymes exist. In a study by Ekstrom et al. (2014), 35 plant genomes were screened for genes encoding putative carbohydrate active enzymes, and corresponding genes were collected in the PlantCAZyme database. To date, this database contains a total of 476 plant-derived AA3 enzymes, most of which are still uncharacterized. As a result, these enzymes have not been assigned to specific subfamilies, and it is not known if the four defined subfamilies that are used for fungal enzymes are applicable to plant enzymes.

Most of the functional data from plant AA3 enzymes are based on reverse genetic studies where the enzyme function has been disrupted and the following phenotypic effects are evaluated. At the functional level, one of the best characterized AA3 family members in plants is HOTHEAD (HTH) from Arabidopsis (*Arabidopsis thaliana*). A mutation in the *HTH* gene causes aberrant flower organ fusions, misshapen epidermal cells, and increased cuticle permeability (Krolikowski et al., 2003; Kurdyukov et al., 2006; Lolle et al., 1998). In addition, the mutant plants have lower amounts of C16 and C18 α-, ω-dicarboxylic fatty acids, and accumulate C16 and C18 ω-hydroxyacids in leaves. The main function of the HTH enzyme seems to be the oxidation of long chain ω-hydroxyacids to α-, ω- dicarboxylic fatty acids for cuticle biosynthesis (Kurdyukov et al., 2006). *In vitro* studies of the Arabidopsis fatty alcohol oxidase FAO3 showed that this enzyme oxidizes 1-dodecanol and 1- hexadecanol (Cheng et al., 2004). In transgenic Arabidopsis plants, FAO3 and FAO4b together were shown to oxidize long chain fatty alcohols, especially C30, to form the corresponding aldehydes (Yang et al., 2022). A corresponding overexpression and knockout plants showed morphological changes in the cuticular waxes of their stems. In *Nesocodon mauritianus*, NmNec3 is involved in color formation of nectar by converting sinapyl alcohol to the sinapyl aldehyde needed for pigment formation (Roy et al., 2022).

In this study, we present the first in-depth biochemical characterization and crystal structure of a plant AA3 family enzyme, which is also the first AA3 enzyme discovered in conifers, the *Pa*AOX1 alcohol oxidase from Norway spruce (*Picea abies* L. Karst.). The phylogenetic analysis distinguishes *Pa*AOX1 from microbial AA3 enzymes as well as from the previously described fatty alcohol oxidases of Arabidopsis and NmNec3 of *N. mauritianus*. We show that the recombinant *Pa*AOX1 displays the most activity toward glycerol. Based on the enzyme activity, its predicted localization, and tissue-specific gene expression, the putative biological role of the enzyme is discussed.

## Materials and methods

### Screening of Norway spruce genome and phylogenetic analysis

Sequences (42) containing Pfam05199: GMC_oxred_C were subtracted from the Norway spruce genome v1.0 (Nystedt et al., 2013). Short fragments were removed, and 20 sequences with more than 300 amino acid residues were analyzed using dbCAN. The 15 sequences recognized the as AA3s were kept for further analysis. A similar process was used to retrieve 3 silver birch (*Betula pendula*) sequences (Salojärvi et al., 2017). Furthermore, 12 Arabidopsis AA3 sequences were obtained from the TAIR database (Berardini et al., 2015), and 2 sequences from *Marchantia polymorpha* and 12 from *Amborella trichopoda* were retrieved from PLAZA Dicots 5.0 (Van Bel et al., 2022) based on orthogroup information. Microbial sequences were chosen so that members from all subfamilies were present, and the alditol oxidase QKN69540.1 from *S. coelicolor* was chosen based on its reported activity converting glycerol into glyceric acid. These sequences and the characterized GMC superfamily enzyme *Nm*Nec3 from *N. mauritianus* were obtained from the NCBI GenBank database (Sayers et al., 2022).

Phylogenetic trees were constructed for plant and microbe sequences, and sequence data were aligned using MAFFT version 7.505 (Katoh & Standley, 2013) with the L-INS-i option. Then, the maximum likelihood phylogenetic trees were constructed using RAxML version 8.2.12 (Stamatakis, 2014) with 1,000 bootstrap replicates.

### Cloning of PaAOX1-encoding cDNA

The *Pa*AOX1 encoding cDNA was cloned from the cDNA library. RNA was isolated from tissue- cultured Norway spruce cells and stored at −80 °C (Laitinen et al., 2017). First-strand cDNA was synthesized with SuperScript™ III reverse transcriptase (Thermo Fisher Scientific, Waltham, MA, USA) using 1 µg of total RNA as a template.

The UTR binding primers *Pa*AOX1_UTR_F and *Pa*AOX1_UTR_R, were used to amplify *Pa*AOX1 cDNA using Phusion High-Fidelity Polymerase (Thermo Fisher Scientific, Waltham, MA). PCR products were ligated to pJET1.2 vector (Thermo Fisher Scientific), transformed to *Escherichia coli* DH5α and the isolated pJET1.2_*Pa*AOX1 plasmid was sequenced. The *Pa*AOX1 coding sequence was submitted to a NCBI GenBank with an accession number PP651606. The *Pa*AOX1 was amplified from pJET1.2_*Pa*AOX1 using the primers *Pa*AOX1pp_F and *Pa*AOX1pp_R and ligated to the pPICZαA expression vector (Thermo Fisher Scientific, Waltham, MA) between the KpnI and XbaI sites, transformed to DH5α, and sequenced. pPICZαA_*Pa*AOX1 was linearized with MssI enzyme and transformed to *Pichia pastoris* wild type strain X-33 (Thermo Fisher Scientific, Waltham, MA) using electroporation. Transformants were selected at 30 °C on YPDS plates (1% yeast extract, 2% peptone, 2% dextrose, 1 M sorbitol, 2% agar, and 100 µg/ml Zeocin™ (InvivoGen, San Diego, CA, UDA). The target protein is produced as a C-terminally His-tagged protein under the methanol-inducible AOX promoter and with *α*-factor secretion signal peptide. All the primers used in this study are listed in Table S2.

### Recombinant expression of PaAOX1 in Pichia pastoris and protein purification

The *P. pastoris* transformant with the most *Pa*AOX1 expression based on small scale expression trials was chosen for the enzyme production in 2 L shake flasks containing 500 ml of BMMY medium (100 mM potassium phosphate buffer, pH 6, 2% (w/v) peptone, 1% (w/v) yeast extract, 1.34% (w/v) yeast nitrogen base, 4 × 10^−5^% (w/v) biotin, and 0.5% (v/v) methanol) for 6 days at 20 °C and 160 rpm. Pre-cultures were grown overnight in BMGY medium in which methanol was replaced by glycerol (15% (v/v)) at 30 °C and 200 rpm. Cells were centrifuged at 1500 x g for 7 min at 4 °C and resuspended in BMMY. Methanol was added to 0.5% (v/v) after 24 h and 1% (v/v) on the following days at 24 h intervals. After the cultivation, cells were removed by centrifugation, the culture supernatant was filtered through a 0.45 μM polyethersulfone (PES) filter, concentrated 15-fold, and buffer-exchanged to 50 mM sodium phosphate buffer, 300 mM NaCl, pH 7.8, using a Vivaflow® 200 laboratory Crossflow cassette (Sartorius AG, Göttingen, Germany). *Pa*AOX1 was purified using Ni^2+^ HisTrap™ Fast Flow immobilized metal ion affinity column (Cytiva, Marlborough, MA, USA) and ÄKTA go™ protein purification system (Cytiva) using 50 mM sodium phosphate buffer, 300 mM NaCl, pH 7.8, and an imidazole gradient of 0 to 1 M in 40 min for elution. Fractions containing *Pa*AOX1 were combined, and the buffer was exchanged to 10 mM HEPES, pH 7, using Vivaspin 20 centrifugal concentrator units (PES, 10 kDa; Sartorius AG). Protein was concentrated to 33 mg/ml and stored in aliquots at −80 °C. 20 mg of the recombinant enzyme could be purified from 1 L of culture medium.

The purified recombinant protein showed a high level of glycosylation (Fig. S2). It was deglycosylated under denaturing conditions using PNGase F (New England Biolabs, Ipswich, MA, USA). The band corresponding to the expected size of *Pa*AOX1 was analyzed using liquid chromatography (LC)-MS in the Meilahti Clinical Proteomics Core Facility at the University of Helsinki. The expressed and purified protein matched the amino acid sequence of *Pa*AOX1, and the identified peptides covered 76% of the protein sequence.

### Enzymatic assays

The *Pa*AOX1 enzyme activity was first screened in multiwell plate assays at 30 °C in 50 mM sodium acetate buffer, pH 5, in a 250 µl reaction volume using cellobiose, anisyl alcohol, methanol, glycerol, and a monosaccharide mixture. The final concentration of each substrate was 10 mM. BQ, DCIP, and molecular oxygen were tested as electron acceptors. The concentration of benzoquinone was 0.2 mM, and reduction of BQ (ε_290_ = 2.24 mM^−1^ cm^−1^) was followed at 290 nm. The DCIP concentration was 1 mM, and the reduction (ε_520_ = 7.8 mM^−1^ cm^−1^) was followed at 520 nm. The formation of H_2_O_2_ was coupled to the oxidation of 1 mM ABTS by adding 1.5 U of horseradish peroxidase (HRP), type I, and the oxidation of ABTS (ε_420_ = 36 mM^−1^ cm^−1^) was followed at 420 nm. All reactions were measured using an Eon plate reader (BioTek, Winooski, VT, USA) and were performed in triplicate.

The pH optimum of *Pa*AOX1 activity on glycerol was screened with ABTS at 30 °C in 50 mM Mcllvaine’s buffer at pH values from 3.0 to 7.5, in 50 mM Tris–HCl buffer at pH values from 8 to 9, and in 50 mM glycine–NaOH buffer at pH values from 9.5 to 10.5. All reactions were performed in triplicate.

The initial activity of *Pa*AOX1 with a list of substrates (Table S1) was tested with the ABTS-coupled reactions. Reactions (250 µl) were performed at 30 °C in 50 mM sodium phosphate buffer, pH 7, 10 mM substrates, 1 mM ABTS, and 6 U/ml HRP. Dodecanol, Hexadecanol, and 16-hydroxyhexadecanoid acid had limited solubility at 10mM concentration. Thus, for these and decanol additional experiments with final substrate concentration ranging from 50-250 µM were tested using both acetone and DMSO as solvents as described in (Takahashi et al., 2016). Glycerol was used as a control in the same reaction conditions. Steady-state kinetic constants of *Pa*AOX1 shown in Table 1 were measured using the same conditions as in the initial activity assays. The velocity of the reaction was plotted against the substrate concentration (8 points for each substrate). The Michaelis–Menten constant (K_M_) was estimated by fitting the data to the Michaelis–Menten equation using Origin 2022 (OriginLab, Northampton, MA, USA). All reactions were performed in triplicate.

**Table 1.**
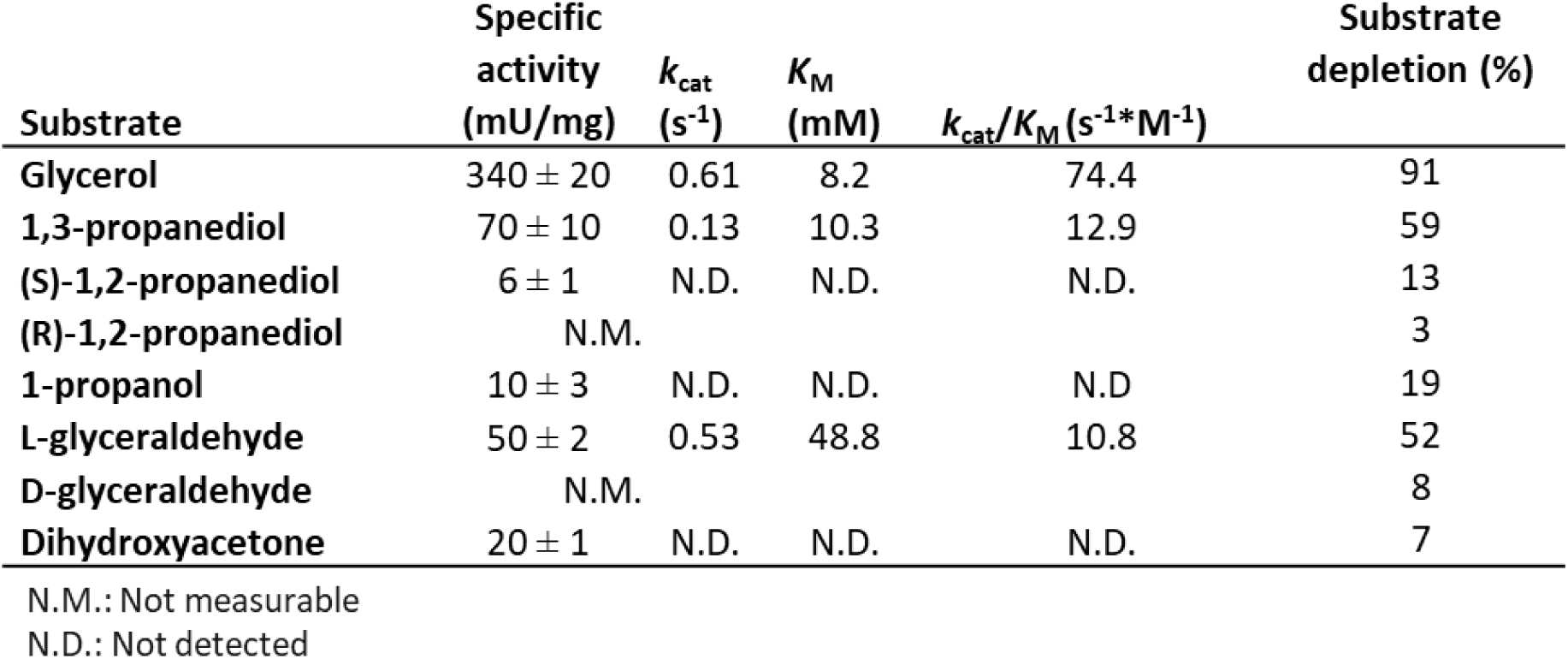
Oxidation of three-carbon substrates by PaAOX1. Specific activity at 10 mM substrate concentration, and kinetic parameters were measured at 30 °C and pH 7. Substrate depletion was determined after 24 h reactions.

| Substrate | Specific activity (mU/mg) | $k_{cat}$ (s<sup>-1</sup>) | $K_M$ (mM) | $k_{cat}/K_M$ (s<sup>-1</sup>*M<sup>-1</sup>) | Substrate depletion (%) |
| --- | --- | --- | --- | --- | --- |
| Glycerol | 340 ± 20 | 0.61 | 8.2 | 74.4 | 91 |
| 1,3-propanediol | 70 ± 10 | 0.13 | 10.3 | 12.9 | 59 |
| (S)-1,2-propanediol | 6 ± 1 | N.D. | N.D. | N.D. | 13 |
| (R)-1,2-propanediol | N.M. |  |  |  | 3 |
| 1-propanol | 10 ± 3 | N.D. | N.D. | N.D. | 19 |
| L-glyceraldehyde | 50 ± 2 | 0.53 | 48.8 | 10.8 | 52 |
| D-glyceraldehyde | N.M. |  |  |  | 8 |
| Dihydroxyacetone | 20 ± 1 | N.D. | N.D. | N.D. | 7 |

N.M.: Not measurable N.D.: Not detected

### Conversion of alcohols by PaAOX1

The conversion of glycerol, D-glyceraldehyde, L-glyceraldehyde, 1,3-propanediol, and 1-propanol were studied. Reactions were performed at 30 °C for 24 h with shaking (450 rpm) in 20 mM sodium phosphate buffer, pH 7, containing 20 mM of substrate and 6 U/ml catalase to remove H_2_O_2_ and recycle part of the oxygen back to the reaction. The *Pa*AOX1 loading was 100 mU/ml (5 mU enzyme per μmole substrate) for each reaction, and control reactions were conducted after deactivating *Pa*AOX1 by boiling for 10 min. The reactions were conducted in duplicate. Time course sampling for the HPLC and MS analysis was done at 3, 9, and 24 h. Reactions were stopped by filtering the samples through 10 kDa cut-off Vivaspin centrifugal columns (Sartorius AG, Göttingen, Germany) to remove the active enzyme.

The filtered samples were analyzed with a LC system (Waters Corporation, Milford, MA, USA) employing a cation exchange Hi-Plex H (300 mm × 6.5 mm, Agilent Technologies, Santa Clara, CA, USA) column. The analytes were monitored with a refractive index (RI) detector. Dihydroxyacetone was monitored with a photodiode array (PDA) detector at 210 nm. The column temperature was 65 °C, and the flow rate 0.5 ml/min. 3 mM sulfuric acid was used as a mobile phase. The quantification of substrate depletion and product formation was done with external standards, including glycerol, glyceraldehyde, dihydroxyacetone, glyceric acid, 1,3-propanediol, and 1-propanol. The standards were prepared in six concentrations to create a standard curve for each run.

The reaction products were analyzed with MS for *Pa*AOX1-oxidized glycerol and 1,3-propanediol. The samples that were incubated with *Pa*AOX1 for 24 h were diluted to 20 µg/ml with 50% methanol, and ammonium chloride was added to a final concentration of 40 µg/ml. MS analysis was performed using quadruple time-of-flight (Q-TOF) MS with an electrospray ionization (ESI) source (SYNAPT G2-Si; Waters Corporation, Milford, MA, USA). The samples were directly infused to the analyzer and analyses were conducted in a negative mode, with the ions collected in a *m/z* range of 50 to 300 at a flow rate of 10 μl/min. The capillary was set to 3 kV; source temperature to 80 °C, and desolvation temperature to 150 °C. The cone gas was 100 l/h and desolvation gas was 600 l/h with nebulizer set at 6 bar.

### Crystallization of PaAOX1 and X-ray data collection

The crystallization of *Pa*AOX1 was performed using the hanging drop vapor diffusion method at 23 °C. For crystallization screening, the Ni-NTA-purified protein was deglycosylated with endoglucanase H (EndoH; DuPont, Wilmington, DE, USA) by adding 1 µg of EndoH for each mg of *Pa*AOX1 and incubating at room temperature for 3 h. Subsequently, the *Pa*AOX1 sample was purified with a Superdex 75 pg size exclusion column (Cytiva, United States) equilibrated with 10 mM HEPES, 0.15 M NaCl buffer, pH 7.0. The purified fractions were pooled and concentrated and buffer- exchanged into 10 mM HEPES, pH 7.0, using Vivaspin 500 centrifugal concentrators (Sartorius AG, Göttingen, Germany) with 10 kDa cut-off membranes. Initial crystallization hit was obtained on a Top96 crystallization screen (anatrace) in conditions containing 0.2 M CaCl_2_, 20 % PEG 3350, with a 1:1 ratio of the crystallization solution and *Pa*AOX1 in 10 mg/ml protein concentration. Crystals obtained in these conditions were used for obtaining an initial *Pa*AOX1 structure, with the data collected at the I24 beamline at the Diamond Light Source synchrotron (DLS) in Oxfordshire, UK. Optimization of crystallization conditions was conducted using native, non-deglycosylated *Pa*AOX1 and yielded the best crystals with a crystallization solution containing 20% PEG 3350, 0.2 M CaCl_2_, and 10 mM glycerol. The crystallization drops were prepared by mixing 1:1 (v/v) the crystallization solution with 33 mg/ml *Pa*AOX1 in 10 mM HEPES buffer, pH 7. Yellow, anisotropic needle crystals suitable for X-ray diffraction analysis were obtained after two weeks’ incubation at 20 °C. Needle crystals were picked with 0.1–0.5 mm loops. Due to high concentrations of PEG 3350 and glycerol, the crystals were not soaked in a cryoprotectant solution prior to being flash frozen in liquid nitrogen. X- ray data for the final *Pa*AOX1 structure deposited into the Protein Data Bank (PDB) were collected at beamline I03 at Diamond Light Source (DLS), using a single *Pa*AOX1 crystal. The beamline was equipped with an Eiger2 XE 16M detector, and the beam was calibrated at a wavelength of 0.953 Å, enabling high-resolution data collection. A total of 3,600 images covering 360° were collected.

### X-ray data processing, structure building, and refinement

For solving the initial *Pa*AOX1 structure, the collected X-ray diffraction data were processed using XDSGUI and the Staraniso server (Brehm et al., 2023; Tickle et al., 2016). Phases were calculated by molecular replacement with Phaser within the CCP4i2 interface, using the PDB structure 1JU2 of almond hydroxynitrile lyase as a search model (McCoy et al., 2007; Potterton et al., 2018). This initial model was refined and built using Refmac and COOT (Emsley et al., 2010; Kovalevskiy et al 2018). The dataset for the final *Pa*AOX1 structure was processed by the automatic data processing pipeline at DLS, utilizing Xia2 software (Evans, 2006; Grosse-Kunstleve et al., 2002), the DIALS package (Winter et al., 2018), and the DIMPLE pipeline (Wojdyr et al., 2013), enabling indexing and integration of the reflection data obtained during automated data collection at Diamond, as well as solving the phases by molecular replacement using the initial *Pa*AOX1 structure as a search model.

The data were processed in the space group P 1 21 1, with an intensity-to-noise ratio of 0.5 at the highest resolution shell (1.64–1.67 Å). The refined unit-cell parameters were defined as: a = 63.1, b = 99.3, and c = 87.9; with α = 90, β = 100.3, and γ = 90. The calculated Vm (Matthew’s coefficient) was determined to be 1.3 Da^−1^ with an estimated two NCS protein molecules in the asymmetric unit. Within CCP4i2, 5% of the data were allotted for R_free_ calculations using _free_ Rflags, and the *Pa*AOX1 structure was further refined with several rounds of iterative model building and refinement using Coot and Refmac5, resulting in a final R_work_ of 0.185 and R_free_ of 0.21 (Emsley et al 2010; Kovalevskiy et al 2018; Potterton et al 2018; Vagin et al., 2004). The statistics of data collection and structure refinement are shown in Table S3. The coordinates for the final *Pa*AOX1 structure model and the structure-factor amplitudes were deposited at the PDB (Berman et al., 2000) with access code 8S2Y. The PyMOL molecular graphics system, version 2.2.3; Schrödinger, Inc was used for graphic visualization of the structure. The volume of the cavity at the catalytic center was calculated using Mole 2.0 software with standard settings (Sehnal et al., 2013).

## Results

### Phylogenetic analysis and expression profiles of PaAOX1 in pre-existing gene expression libraries

The Norway spruce genome version 1.0 (Nystedt et al., 2013) has 42 putative AA3 family genes, 15 of which were predicted by the web server for the automated Carbohydrate-active enzyme ANnotation (dbCAN) to encode AA3 enzymes (Zhang et al., 2018). Phylogenetic analysis was performed to compare these 15 sequences, along with 27 additional annotated plant AA3 sequences with fungal AA3 sequences and the glycerol-oxidizing AldO sequence from *Streptomyces coelicolor*. The analysis showed that microbial sequences clustered together based on their subfamilies (Fig. 1). AldO (QKN69540.1) from *S. coelicolor* formed its own branch, and although it has been reported to be able to convert glycerol into glyceric acid, it did not cluster close to *Pa*AOX1. Most of the plant sequences were phylogenetically distant from the microbial sequences and formed one main plant clade that divided into subclades. However, one exception was a branch of the plant AA3 sequences that included the fatty alcohol oxidases FAO3 and FAO4b. This branch was closer to that containing microbial pyranose oxidase sequences than to other plant sequences in the main plant clade, which suggests a more distant evolutionary origin for these enzymes. The Norway spruce AA3 sequences (PaAA3s) fell mainly into two clusters, one cluster with the known fatty alcohol oxidases, and a second cluster with uncharacterized enzymes. The latter cluster included *Pa*AOX1 (MA_10434427g0010), which was selected for further studies.

**Figure 1.**
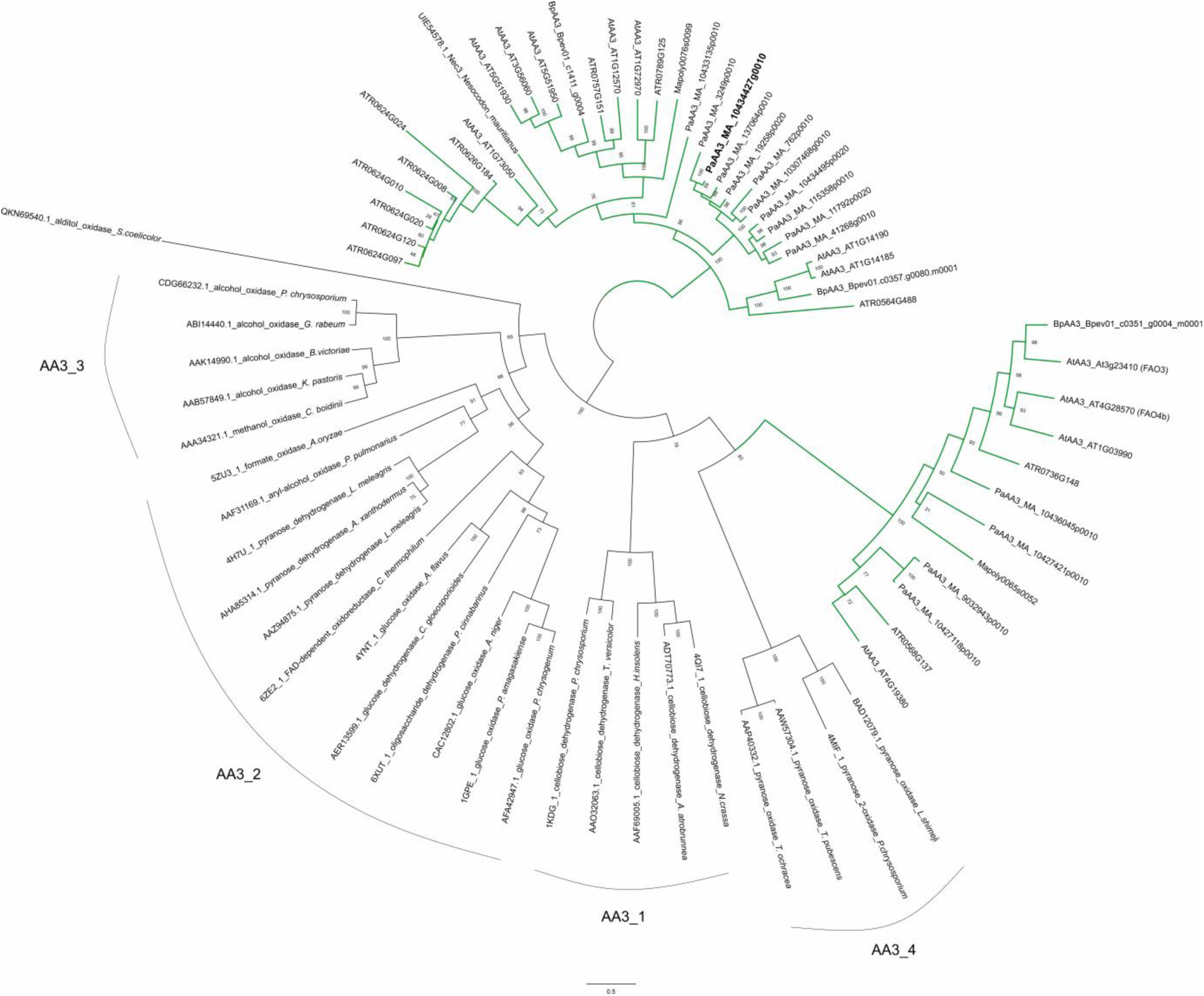
Unrooted maximum likelihood phylogenetic tree was used to compare relatedness of microbial and plant GMC oxidoreductases. Most of the microbial and plant sequences formed their own clades. For microbial sequences, a clear separation based on AA3 subfamilies was observed. Some plant enzymes, including fatty alcohol oxidases FAO3 and FAO4b, clustered with the fungal AA3_4 subfamily pyranose oxidases instead of other plant sequences forming the main plant clade. PaAOX1 is in bold and localized within the main plant clade. None of the other enzymes in the same branch have been characterized. Branches containing plant sequences are marked with a green color. Bootstrap values are based on 1,000 replicates.

The expression pattern of the *PaAOX1* gene was searched from the available Norway spruce RNA- sequencing expression atlases (Sundell et al., 2015). In the database, the expression of *PaAOX1* was highest in needles and vegetative shoots. Lower level of expression was present in the wood sample harvested in August that contained phloem, cambium, and xylem (Fig. S1). The higher resolution gene expression resource NorWood did not contain *PaAOX1* transcripts (Jokipii-Lukkari et al., 2017).

### Characterization of PaAOX1 activity

The *PaAOX1* gene encodes a protein with a predicted molecular weight of 61 kDa. The protein has a signal peptide of 23 amino acids, and it is predicted to be localized in the apoplast. Initially, the *Pa*AOX1 enzyme activity was screened at pH 5 using different electron donors, including cellobiose, anisyl alcohol, methanol, glycerol, and a monosaccharide mixture containing D-glucose, D-mannose, L- arabinose, D-xylose, and D-galactose. Benzoquinone (BQ), 2,6-dichlorophenolindophenol (DCIP) and molecular oxygen were tested as electron acceptors. Under these conditions, *Pa*AOX1 showed activity toward glycerol. Only oxygen functioned as an electron acceptor, and thus, H_2_O_2_ was concomitantly produced. The optimum activity of *Pa*AOX1 was detected to be at pH 7, and the enzyme maintained at least 80% of its activity between pH 4 and 8 (Fig. S3). A broader screen of *Pa*AOX1 at the optimal pH included 54 electron donors selected based on previously reported AA3 activities and plant phenotyping studies (Table S1). The analysis revealed that *Pa*AOX1 was primarily active to substrates containing three carbons (Fig. 2). The highest specific activity in decreasing order was on glycerol, 1,3- propanediol, L-glyceraldehyde, dihydroxyacetone, 1-propanol, and (S)-1,2-propanediol (Table 1).

**Figure 2.**
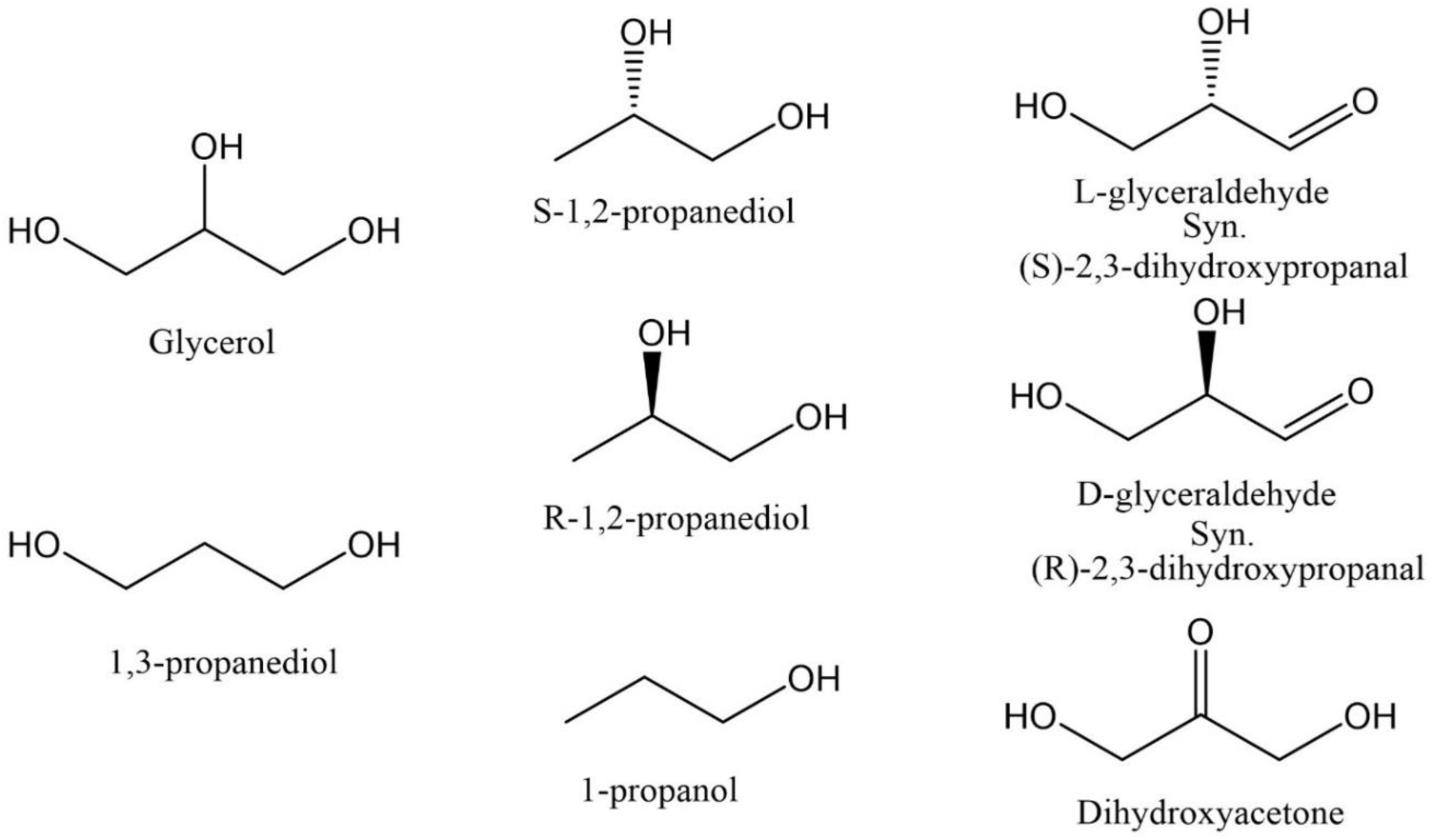
PaAOX1 showed catalytic activity toward electron donors containing three carbons.

*Pa*AOX1 displayed the highest catalytic efficiency with glycerol, and 91% of glycerol conversion was achieved within 24 h at 30 °C and pH 7. *Pa*AOX1 demonstrated comparable catalytic efficiency for both 1,3-propanediol and L-glyceraldehyde, but greater affinity to 1,3-propanediol and higher turnover rate for L-glyceraldehyde. *Pa*AOX1 showed clear enantioselectivity. It oxidized both L-glyceraldehyde and (S)-1,2-propanediol, whereas no oxidation was detected in the plate assays with D-glyceraldehyde or (R)-1,2-propanediol. The enantioselectivity of *Pa*AOX1 was further confirmed with high- performance liquid chromatography (HPLC). Conversion of 52% of L-glyceraldehyde was measured at 24 h in comparison to conversion of 7.6% of D-glyceraldehyde, and 13.0% (S)-1,2-propanediol was compared to 2.5% (R)-1,2-propanediol at 24 h (Table 1).

Oxidation by *Pa*AOX1 of 20 mM of glycerol and L- and D-glyceraldehyde was followed using HPLC for product identification and quantification (Fig. 3). The primary product of glycerol was glyceraldehyde, the concentration of which reached 10 mM after 9 h (Fig. 3A). Glyceraldehyde can react with water to generate a hydrate, geminaldiol, and *Pa*AOX1 was able to further oxidize the hydrate of glyceraldehyde to glyceric acid. The formation of glyceric acid was detectable at 9 h and reached 5 mM after 24 h. Additionally, a small amount of dihydroxyacetone was formed. The oxidation of glycerol by *Pa*AOX1 was also followed using mass spectrometry (MS) to detect products beyond glyceraldehyde, dihydroxyacetone, and glyceric acid. However, no additional products were detected (data not shown). In the reaction with L-glyceraldehyde (Fig. 3B), *Pa*AOX1 oxidized the hydrate form of L-glyceraldehyde primarily to glyceric acid, while a minor amount of dihydroxyacetone was also detected. D-glyceraldehyde was poorly oxidized by *Pa*AOX1 (Fig. 3C), with only 0.2 mM and 0.5 mM of glyceric acid and dihydroxyacetone detected after 24 h, respectively. Notably, a small amount of glycerol (1 mM) was detected as an impurity in both the L-glyceraldehyde and D-glyceraldehyde used in this study (Fig. 3B, C). None of the products were detected in the control reactions comprising heat inactivated *Pa*AOX1 (Fig. S4).

**Figure 3.**
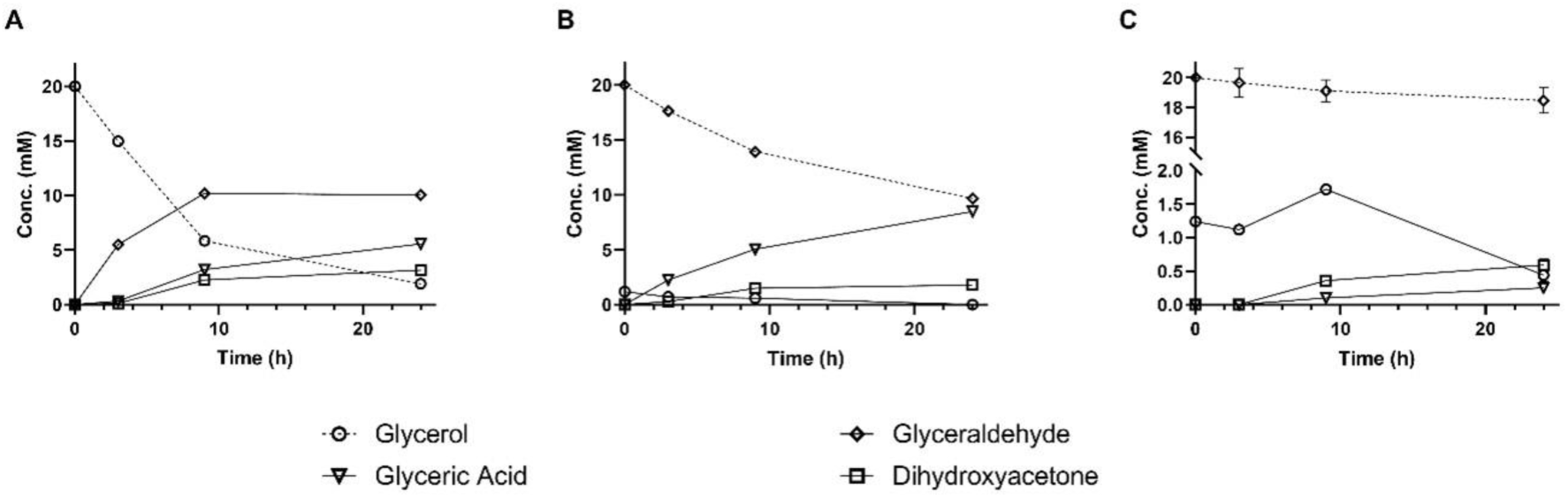
Product formation and substrate depletion catalyzed by PaAOX1 was followed for 24 h using HPLC. The depletion of the substrates is indicated with dashed lines, and the formation of products is shown with solid lines. Error bars show the standard deviation of replicate samples (n = 3). A) Glycerol was used as a substrate. After 24 h, approx. 10 mM glyceraldehyde, 5 mM glyceric acid, and 2.5 mM dihydroxyacetone were formed. There was 2.5 mM of glycerol still remaining after 24 h incubation. B) When L-glyceraldehyde was used as a substrate, approx. 10 mM of the substrate was converted to 10 mM of glyceric acid. Trace amounts of dihydroxyacetone were detected. C) When D-glyceraldehyde was used as a substrate, approx. 7.6% of the substrate depletion was detected after 24 h. Minute amounts of glyceric acid and dihydroxyacetone were detected in B) and C), because the D- and L-glyceraldehyde that were used as substrates had glycerol as an impurity.

### Protein structure determination and model building

The crystal structure of *Pa*AOX1 was solved by X-ray crystallography and refined at 1.6 Å resolution, with final R and R_free_ values of 0.18 and 0.21 (Fig. 4). The structure consists of 1,096 amino acids, 762 water molecules, 11 carbohydrate residues, 1 glycerol molecule, 4 sodium ion, and 1 calcium ion. The crystallization solution had 10 mM glycerol, and glycerol molecule was found being bound to *Pa*AOX1 but not at the catalytic center of the enzyme.

**Figure 4.**
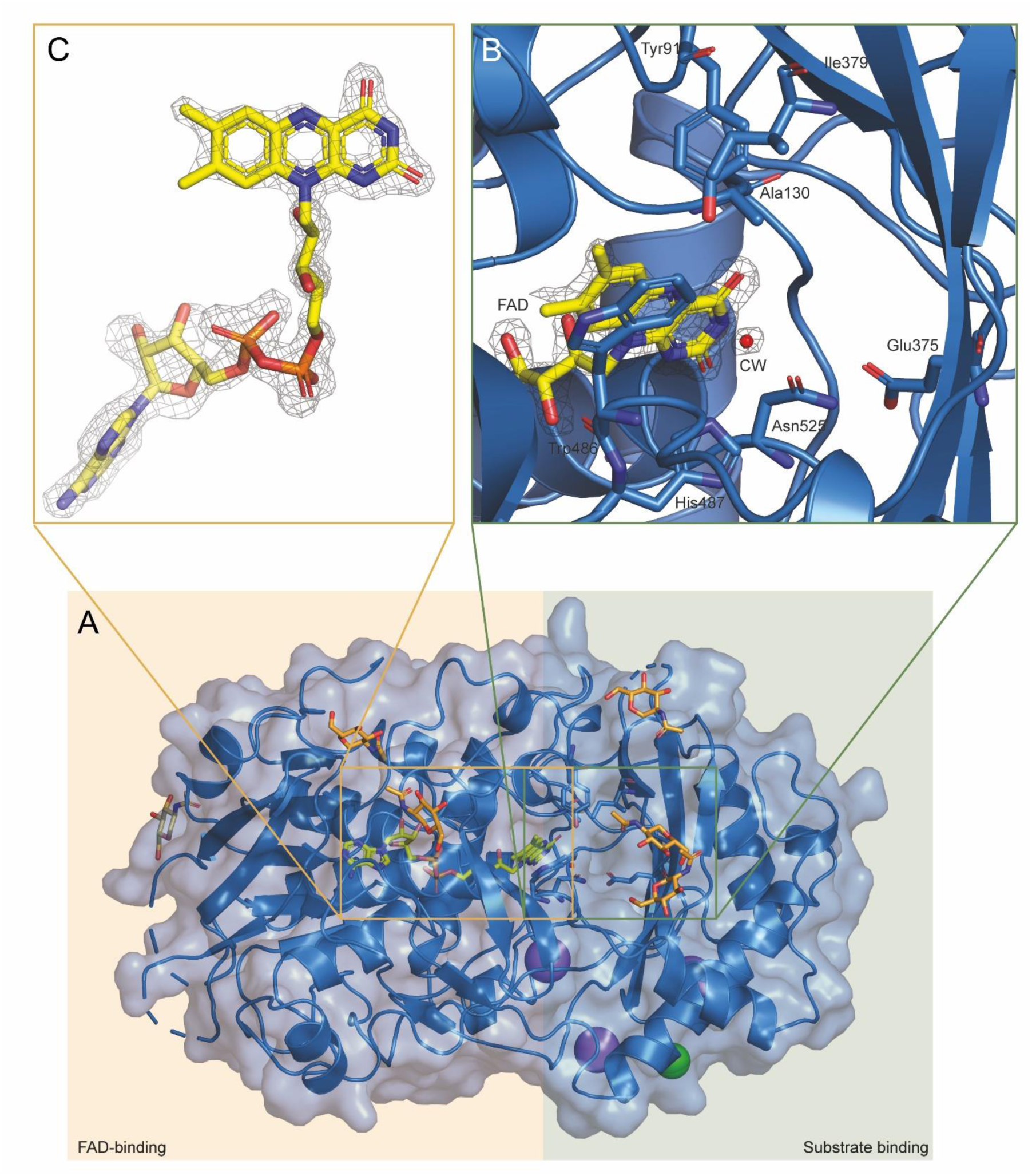
A) Crystal structure of PaAOX1 consists of two domains: a FAD-binding domain and a substrate-binding domain. The crystal structure of PaAOX1, determined to a resolution of 1.64 Å, is depicted. The figure highlights key structural features of an alcohol oxidase, including glycosylations (orange) and the non-covalently bound FAD cofactor (yellow). Additionally, the figure identifies a magnesium ion (green) and additional glycerols, while dotted loops represent regions with missing density. B) In the upper right panel, the active site of PaAOX1 is presented, featuring substrate- interacting amino acids (shown in sticks and labeled), the FAD cofactor, and the conserved water molecule (CW, W1003), in red). C) These elements are superimposed with the final electron density map (gray mesh, 1.0 σ), specifically showing the density of the FAD.

The crystal structure of *Pa*AOX1 has two non-crystallographic symmetry (NCS)-related protein molecules, A and B, in the asymmetric unit, with an average root mean square deviation (RMSD) of 0.134 Å between the two NCS molecules, including all atoms except water. This RMSD value indicates that the two NCS molecules are practically identical. Individual corresponding amino acids in the two NCS models do show different conformations, but these differences are not close to the catalytic center and should therefore not have any significant effect on the catalytic activity of the enzyme.

Within each of the two NCS molecules of the *Pa*AOX1 structure, three surface loops of 242–254; 276– 296 and 383–390 lack a visible electron density, suggesting high flexibility (Fig. S5). The remaining amino acids in the structure display well-ordered electron density, apart from a surface turn (463–465) with a less well-defined electron density in molecule A and could not be built in molecule B (Fig. S5).

The N-terminal of *Pa*AOX1 forms a FAD-binding module that has the classical alpha/beta fold of this class of enzymes. The FAD molecule is non-covalently bound in the catalytic center of each of the two NCS *Pa*AOX1 molecules in the structure. The density of the FAD molecule bound in the catalytic center is a common feature in all substrate-binding domains of the characterized GMC family members (Cavener, 1992). The density in both molecules shows a so-called butterfly FAD (Walsh & Miller, 2003). As no data were obtained to prove a butterfly or planar FAD, the FAD was modeled into the density acquired by the data. This effect can be due to crystallization conditions, oxidation states, and different conformations of the FAD in the lattice of the proteins. The substrate-binding domain of *Pa*AOX1 consists of eight sheets and eight helices built up mainly by the C-terminal of the protein, contributing to its specific structural arrangement (Fig. 5).

**Figure 5.**
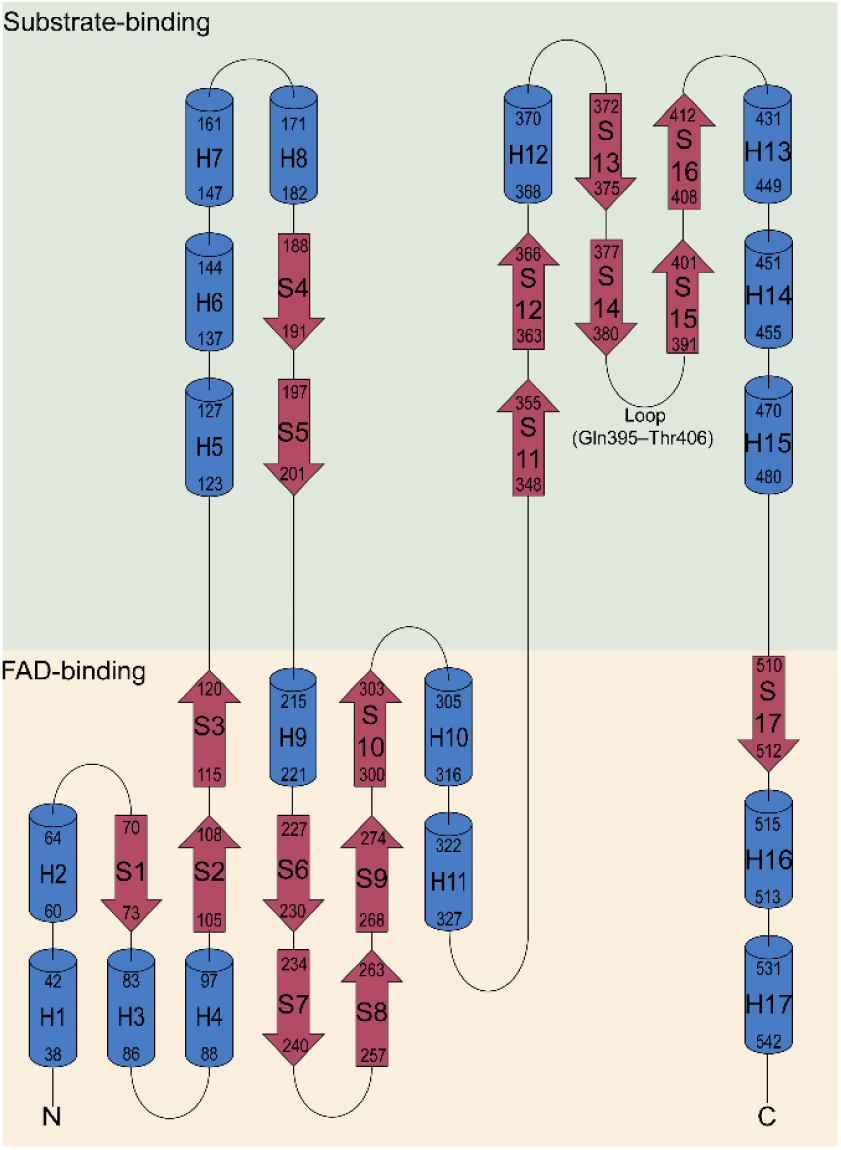
Topology diagram of PaAOX1. This figure illustrates the secondary structural organization of flavin adenine dinucleotide (FAD) and substrate-binding domains within the crystal structure of alcohol oxidase PaAOX1. The helices (in blue) and sheets (in red) that are part of the FAD- and substrate-binding domains are shaded in yellow and green, respectively. The start and end of the helices and sheets are indicated by amino acid numbers, corresponding to the sequence alignment and the structure. Additionally, a loop presumed to play a role in the arrangement of the active site and substrate binding is indicated, spanning between sheets 14 and 15.

N-Glycosylation of *Pa*AOX1 is observed at seven asparagine residues in both NCS molecules A and B at positions Asn100, Asn232, and Asn382. An additional N-glycosylation is found at Asn275 in NCS molecule B. Notably, the glycosylation at Asn382 is close to the entrance of the active site, at the end of the beta-sheet C3 and at the beginning of the loop consisting of residues 383–390, which is flexible in structure and presumably forms a loop that is part of the substrate binding domain (Fig. 5). The 383– 390 loop lacks density for six amino acids, making it impossible to determine its exact conformation.

However, the beta-sheets C3 and C4 clearly tilt upward, and the beginning of the loop, between these two sheets, does not extend toward the active site (Fig. 4 and 5). As a result, the loop does not cover the entrance of the active site, and thus creates an open tunnel conformation towards the catalytic center.

The absence of density and the presence of glycosylation suggest that the 383–390 loop is flexible and can adopt various conformations and this flexibility may lead to an open and closed active site state.

The catalytic center of GMC family oxidases is located near the FAD isoalloxazine ring (Fernández et al., 2009), where the C-terminal substrate-binding domain of the enzyme has a large cavity formed by the surrounding loops and a four-stranded beta-sheet (Fig. 4). The solvent-accessible cavity at the catalytic center of *Pa*AOX1 was measured using Mole 2.0 software (Sehnal et al., 2013). The cavity has an approximate width of 12 Å and an approximate length of 13.3 Å, measured from Ser483 to FAD. The approximate volume of the catalytic center cavity is 550 Å^3^.

Two amino acids are important for the two half-reactions of the enzyme. One is a strictly conserved histidine residue among AA3 enzymes (His487 in *Pa*AOX1), which is used as a catalytic base taking a proton from the hydroxyl group of the substrate (Romero & Gadda, 2014). The other residue is, in most cases, a histidine, but in *Pa*AOX1 and a few other AA3 enzymes, such as PcAOX from Phanerodontia chrysosporium (Nguyen et al., 2018) and PpAOX1 from *Phanerochaete chrysosporium* (Koch et al., 2016), this histidine is replaced by an asparagine residue (Asn525 in *Pa*AOX1), which presumably acts as a hydrogen donor in the catalytic reaction (Figs. 4, 6). Additional catalytic center residues found in *Pa*AOX1 are Trp486, Ala130, Ser396, and Glu75. In other GMC family members with a known structure, the amino acids at these positions often have aromatic properties and extend into the active center. Small amino acids such as alanine, as in *Pa*AOX1, are relatively rare.

*Pa*AOX1 has the most structural similarity to the almond hydroxynitrile lyases *Pa*HNL1 and *Pa*HNL5, with RMSD values of 0.61, although their catalytic activity is different from *Pa*AOX1 (Dreveny et al., 2001; Pavkov-Keller et al., 2016). *Pa*HNLs have evolved from the GMC family of aryl alcohol oxidases and lost the oxidative activity during evolution (Dreveny et al., 2001). Based on the sequence similarity analysis and RMSD calculations, other homologues in order of most to least similar, are the aryl alcohol oxidase *Pe*AAO from *Pleurotus eryngii* (RMSD: 1.57; Carro et al., 2017), the alcohol oxidase *Pc*AOX (RMSD: 2.08; Nguyen et al., 2018), and the cholesterol oxidase ChOx from *Streptomyces sp*. (RMSD: 8.277; Golden et al., 2017). The overlaid structures and comparison of the active sites of *Pa*AOX1, *Pa*HNL, *Pe*AAO, and *Pc*AOX are shown in Fig. 6 A, B, and C, respectively.

**Figure 6.**
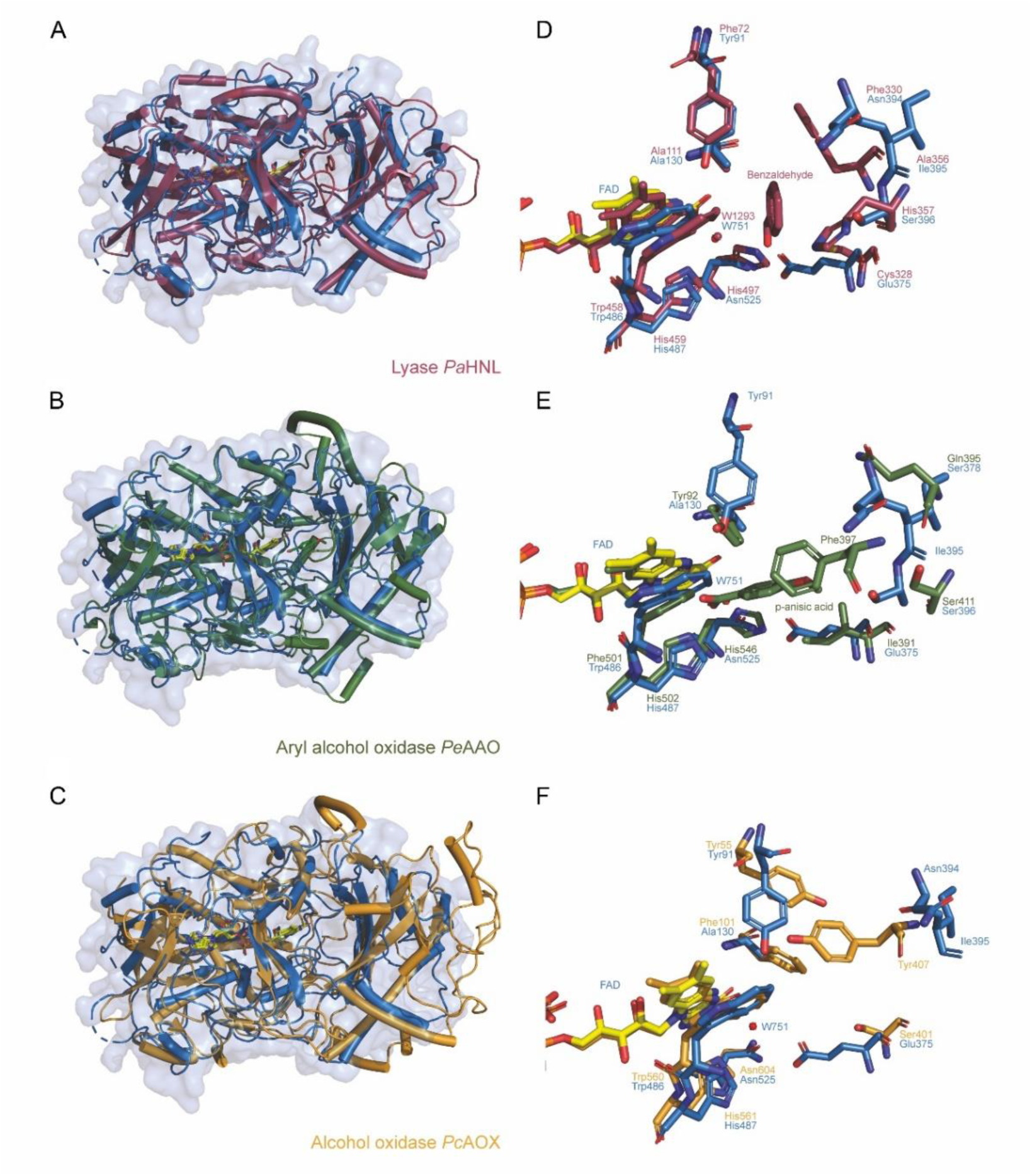
Comparison of the PaAOX1 structure with GMC oxidoreductases with the structural homologues. **A–C:** the crystal structure of PaAOX1 (blue) is compared with its structural homologues: A) The hydroxynitrile lyase PaHNL from almond (Prunus amygdalus) (PDB: 3GDN) in red. B) The aryl alcohol oxidase PeAAO from Pleurotus eryngii (PDB: 5OC1) in green. C) The alcohol oxidase PcAOX from Phanerochaete chrysosporium (PDB: 6H3G) in orange. A structural alignment of cartoon representations of PaAOX1 with the corresponding enzyme structures is displayed, conducted by aligning the FAD in both enzyme structures, thus highlighting the differences in the overall fold among these enzymes when compared to PaAOX1, especially with regard to structural arrangement of the loops in front and those forming the active site. **D–F**: A detailed comparison of the active site of PaAOX1 with those in its structural homologues is presented, showcasing proposed substrate-interacting amino acids (shown as sticks and labeled), the FAD cofactor, glycerol (yellow), and the conserved water molecule (W751, in red). These elements are superimposed with the corresponding active site features of D) PaHNL, E) PeAAO, and F) PpAOX1, revealing distinctions in the active site architecture of these enzymes. PaHNL harbors its inhibitor, benzaldehyde, at a position analogous to PaAOX1’s substrate glycerol (904). Furthermore, PaHNL contains a conserved water molecule (W1293) in proximity to the equivalent W1003. In PeAAO, the substrate p-anisic acid assumes a distinct position close to the isoalloxazine ring of the FAD.

The amino acid sequence comparisons are shown in Fig. 7. The sequence alignment highlights the low amino acid sequence identity of these structural homologues. Moreover, the overall structural similarity as indicated by the RMSD values does not necessarily imply functional similarity due to the different activities of members of the AA3 family, despite having a similar overall fold.

**Figure 7.**
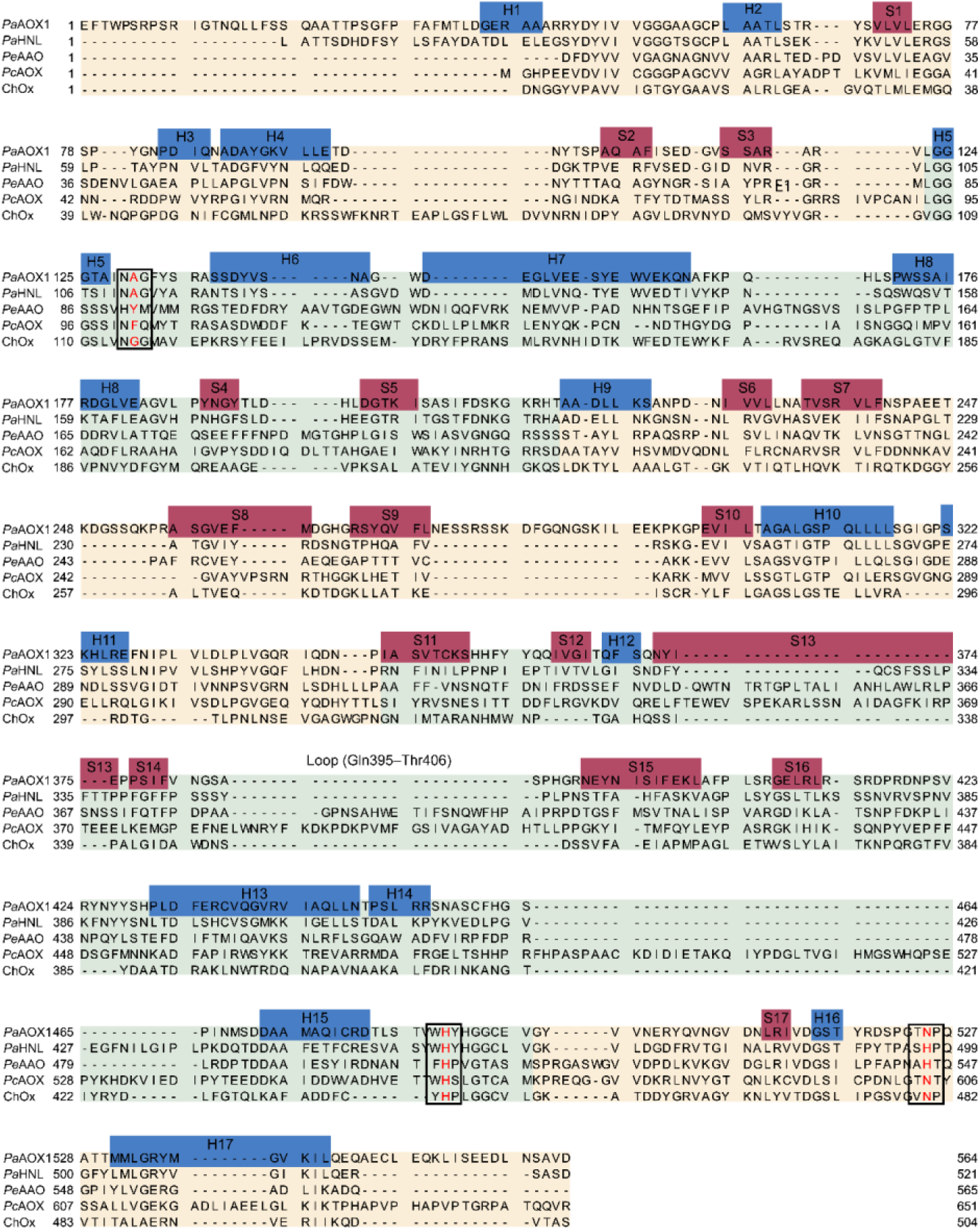
Sequence alignment illustrating the comparison between alcohol oxidase PaAOX1 and its structural homologues: hydroxynitrile lyase PaHNL from almond (Prunus amygdalus), aryl alcohol oxidase PeAAO from Pleurotus eryngii, alcohol oxidase PpAOX1 from Phanerochaete chrysosporium, and cholesterol oxidase ChOx from Streptomyces sp. The multiple sequence alignment was generated using MUSCLE (MUltiple Sequence Comparison by Log Expectation). In the alignment, PaAOX1’s FAD-binding and substrate-binding domains are colored in yellow and green, respectively. Helices are marked in blue and sheets in red with numbering. A black box highlights the proposed active site amino acids, Ala130, His487, and Asn525, along with the corresponding amino acids at these positions in the structural homologues. These amino acids are colored in red to emphasize variations among the proteins. Additionally, a loop presumed to play a role in the active site arrangement and substrate binding is indicated, spanning from the amino acid Gln395 to Thr406. Accession numbers for the proteins used in this alignment are as follows: PaAOX1 (PP651606), PaHNL (AF412329), PeAAO (AAC72747), PpAOX1 (HG425201), ChOx (M31939 J03356).

## Discussion

### Biological role of PaAOX1 based on the enzyme activity and tissue-specific gene expression

Plants have a variety of AA3 oxidoreductases but only a few of them have been biochemically or functionally characterized. The main *in vitro* activity detected from the newly discovered Norway spruce AA3 enzyme, *Pa*AOX1, was the oxidation of glycerol into glyceraldehyde and, further, into glyceric acid. Similar activity has never been reported for any plant species, and based on the phylogenetic analysis, *Pa*AOX1 is positioned in a cluster where none of the enzymes have been functionally characterized. In fungi, only a few other glycerol-oxidizing enzymes have been discovered and characterized (Cleveland et al., 2021; Lin et al., 1996; Linke et al., 2014; Mollerup et al., 2019; Uwajimas et al., 1984). Most are reported to oxidize glycerol only to D-glyceraldehyde except for the homologous *Fusarium graminearum* and *F. oxysporum* enzymes, *Fgr*AAO and *Fox*AAO, which convert glycerol to predominantly L-glyceraldehyde. But the formation of glyceric acid has not been reported (Cleveland et al., 2021). Bacterial alditol oxidase from *S. coelicolor* can convert glycerol into D-glyceric acid with low catalytic efficiency (Gerstenbruch et al., 2012; Heuts et al., 2007).

According to the sequence information, *Pa*AOX1 contains a signal peptide and is secreted into the apoplast. Based on the detected enzymatic activity of *Pa*AOX1 and the tissue-specific gene expression, the biological role can only be hypothesized. Glycerol exists in plants as a free form, but high concentrations are toxic (Eastmond, 2004; Gerber et al., 1988; Wei et al., 2004). Glycerol is also present in the apoplast, where it is consumed, for instance, during plant–pathogen interactions and accumulates during salt stress (Franzisky et al., 2023; O’Leary et al., 2016; Wei et al., 2004). No data on the presence of glyceraldehyde or glyceric acid or information about their importance in the apoplast were found.

In addition to its free form, glycerol is a major constituent of the heteropolymers cutin and suberin. Cuticle is formed on the outer surface of the epidermal cell walls in the aerial parts of plants, and suberin is deposited on the inner surface or inside of the primary cell wall in the bark and in the root epidermis and exodermis, as well as in parts of the endodermis (Beisson et al., 2012). In addition, it has been recently shown that suberin-like components are present in mature wood cell walls in both hardwood and softwood species, including Norway spruce (Derba-Maceluch et al., 2023). In the available Norway spruce transcription data, *Pa*AOX1 is expressed in different tissues, such as needles, which have thick cuticle, vegetative shoots, and in the wood sample (Fig. S1). No transcription data is available from roots or bark. The current evidence proposes that in both cutin and suberin, the long chain α-hydroxyacids and α,ω-diacids are linked via ester bonds to hydroxyl groups of glycerol mediated by acyltransferases (Beisson et al., 2012; Graça, 2015). The possibility of glyceric acid or glyceraldehyde being linked to hydroxyacids has not been reported. It has been suggested that apoplastic H_2_O_2_ would be needed in the polymerization of the dark lamellae structure of suberin (Beisson et al., 2012; Serra & Geldner, 2022).

It is possible that the main function of *Pa*AOX1 is the production of H_2_O_2_ in the apoplast, and thus, H_2_O_2_ would be the biologically important molecule instead of the oxidized product, such as glyceraldehyde or glyceric acid. It is well known that H_2_O_2_ has many roles in plants, from signaling to lignification, and an apoplastic reactive oxygen species (ROS) burst is an important plant defense response (Bolwell et al., 2002; Considine & Foyer, 2021; Kärkönen & Kuchitsu, 2015). Transgenic plants would be needed to resolve whether glycerol is the main substrate of *Pa*AOX1 *in planta*.

### Structural features of *Pa*AOX1 in comparison to other GMC superfamily members

Based on the structure, *Pa*AOX1 belongs to the GMC oxidoreductase superfamily, which consists of two domains typical for the family: the N-terminal FAD-binding domain and the C-terminal substrate- binding domain (Cavener, 1992). The major structural difference between *Pa*AOX1 and its structural homologues is in the organization of the substrate binding domain with a loop ranging from residues 383–390, close to the active site. In *Pa*AOX1, the 383–390 loop between sheets C3 and C4 (Fig. 4) bends upward and points away from the active site. The loop is also present in the two microbial AA3 alcohol oxidases *Pc*AOX and *Pp*AOX. The active sites of *Pc*AOX and *Pp*AOX are solvent- inaccessible but not covered by the loop (Fig. 6C; Koch et al. 2016; Nguyen et al., 2018). In *Pa*HNL, the loop covers the active site, and it is proposed that the substrate enters through a 16 Å long tunnel above the helices H3 and H4 (Fig. 6A) (Dreveny et al., 2001; Pavkov-Keller et al., 2016). In *Pe*AAO, the substrate *p*-anisic acid is only accessible through a small hydrophobic funnel (Fernández et al., 2009). However, molecular dynamic simulations indicate that the loop in *Pe*AAO is flexible enough to allow the substrate entrance (Hernández-Ortega et al., 2011).

AA3 structures resolved with and without substrates do not show reorganization of the active site (Carro et al., 2017). Therefore, the importance of loops and reorganization of the active site for substrate binding remains unknown. However, some ChOx enzymes show structures with both open and closed conformations of the active site and are proposed to control the entrance of ligands into the binding pocket, indicating a regulatory function of the loop (Li et al., 1993). The absence of electron density and presence of glycosylation in the loop region surrounding the catalytic center of *Pa*AOX1 indicates that the loop region is flexible and may have a role in controlling the substrate binding in the catalytic center of *Pa*AOX1.

### Active site amino acids and their impact on AA3 enzyme activity

The structures of *Pp*AOX and *Pe*AAO show that the space for the substrate molecules near the active site is limited by the presence of large aromatic amino acids (Carro et al., 2017; Ferreira et al., 2015; Koch et al., 2016). Corresponding residues in the structure of *Pa*AOX1 are Ala130, Ile395, and Ser396. The side chain in Trp486, which is situated approximately 4 Å away from the FAD, is conserved among alcohol and aryl alcohol oxidases in the AA3 family (Carro et al., 2017; Koch et al., 2016). This hydrophobic amino acid appears to be important for FAD binding and may additionally restrict binding of large substrates in the catalytic center of the enzyme.

Exploration of the mutant variants of *Pe*AAO revealed the catalytic significance of specific amino acids within AAO enzymes. In *Pe*AAO, the residue Tyr92 is known to be important for substrate interaction and alcohol stabilization at the active site of aryl alcohol oxidase and in the AA3 family this position is commonly occupied by aromatic and large amino acids such as phenylalanine or tyrosine (Ferreira et al., 2006; Hernández-Ortega et al., 2011; Koch et al., 2016; Nguyen et al., 2018). In the *Pa*AOX1 structure, the corresponding amino acid is Ala130. The loss of aromatic residue at this position led to a reduced specific activity toward aryl alcohols (Kadowaki et al., 2020; Zhao et al., 2024). In *Pe*AAO, mutation of tyrosine to alanine (Tyr92Ala), a mutation that matches the corresponding amino acid in *Pa*AOX1, converted *Pe*AAO to inactive, supporting the involvement of π–π interactions of active site amino acids and aryl-alcohol catalysis in AAO enzymes (Ferreira et al., 2015). Additionally, the residue Glu389 in *Pe*AAO appears to be vital for forming a protonated glutamic acid residue following spontaneous proton transfer from the catalytic His546. In *Pa*AOX1, however, His546 is replaced by Asn525, an observation suggesting a different proton transfer mechanism in the active site and raising questions about the possibility of charge separation, as observed in *Pe*AAO (Hernández-Ortega et al., 2011).

In *Pc*AOX, the mutant with Phe101Ser showed an important change in catalytic activity. Mutation of this phenylalanine to a short amino acid such as a serine, which corresponds to Ala130 in *Pa*AOX1, increased the activity toward glycerol (Nguyen et al., 2018). It appears that a more spacious binding site and possible hydrogen bonding to the substrate are important for glycerol binding, and the presence of serine at this position in *Pc*AOX increased the enzyme activity (Koch et al., 2016; Nguyen et al., 2018). Based on these data, the amino acids Asn525 and Ala130 at the active site of *Pa*AOX1 are likely to have a major influence on the catalytic activity of the enzyme.

A conserved water molecule (CW, W1003) in *Pa*AOX1 is found between N5 of the isoalloxazine ring and the imidazole of the conserved histidine (Fig. 4). The water molecule is present at a similar position in many structures of the AA3 GMC oxidoreductase family, such as in the lyase *Pa*HNL (Dreveny et al., 2001), aryl-alcohol oxidase *Pe*AAO (Carro et al., 2017), alcohol oxidase *Pp*AOX1 (Koch et al., 2016), and cholesterol oxidase ChOx (Fig. 6 A, C; Yue et al., 1999). In *Pe*AAO, the oxygen (O1) in the substrate *p*-anisic acid and the conserved water in *Pa*AOX1 (Fig. 6B) are at nearly the same position. This observation suggests that the conserved water molecule gets replaced by the substrate at the catalytic center, and the catalytic reaction mechanism can take place (Meyer et al., 1998; Wohlfahrt et al., 1999).

Our findings on *Pa*AOX1, the first coniferous AA3 family enzyme to be characterized in depth for biochemical properties and structure, significantly contribute to a deeper understanding of the structural and functional aspects of enzymes in the GMC superfamily. To date, the CAZy database has contained the structures of only fungal alcohol oxidases. The characterized *Pa*AOX1 has a novel activity among the characterized AA3s in preferring the L-glyceraldehyde as a substrate. Information about the protein structure can guide engineering of *Pa*AOX1 for improved catalytic efficiency and adds to understanding of versatile AA3 oxidoreductases in general given that only a limited number of structures have been solved.

## Acknowledgment

The authors would like to thank Diamond Light Source for beamtime (Uppsala BAG) and the staff of beamlines I02, I03, and I24 for assistance with crystal testing and data collection. This work was supported by the Research Council of Finland (decision 308998 to AK, decision 308997 to MT, decision 308996 to EM, and decision 331853 to TP) by Novo Nordisk Foundation (decision NNF21OC0070816 to NS, JK, and MS).

## Author Contributions

TP and NS contributed equally to this work. TP performed cloning, recombinant protein production and purification, part of enzymatic assays and data analyses. NS and TH performed protein crystallization and building of the protein structure. HZ performed part of the enzymatic assays and conducted HPLC and MS analyses and related data analyses. MS and JK performed part of the recombinant protein production, purification, and enzymatic assays. AV generated the phylogenetic tree. TP and NS drafted the manuscript. ERM and MaS contributed to the data interpretation and in the writing of the final manuscript. MT, AK and TP conceived and coordinated the study. All authors read and approved the final manuscript.

## Supporting information

Fig.S1 The PaAOX1 expression profile in Plantgenie.org database.

Fig. S2 Deglycosylation of the purified recombinant *Pa*AOX1

Fig.S3 Optimization of pH for *Pa*AOX1 assays

Fig.S4 Control reactions with boiled *Pa*AOX1

Fig.S5 Loops of PaAOX1 with missing density

Table S1 Compounds tested as substrates

Table S2 Primers used in this study

Table S3 Diffraction data and refinement statistics for the PaAOX1 structure

