## Supporting information for "A glycerol oxidase from Norway spruce (*Picea abies* L. Karst.) expands biochemical and structural attributes of the GMC oxidoreductase superfamily"

The following Supporting Information is available for this article:

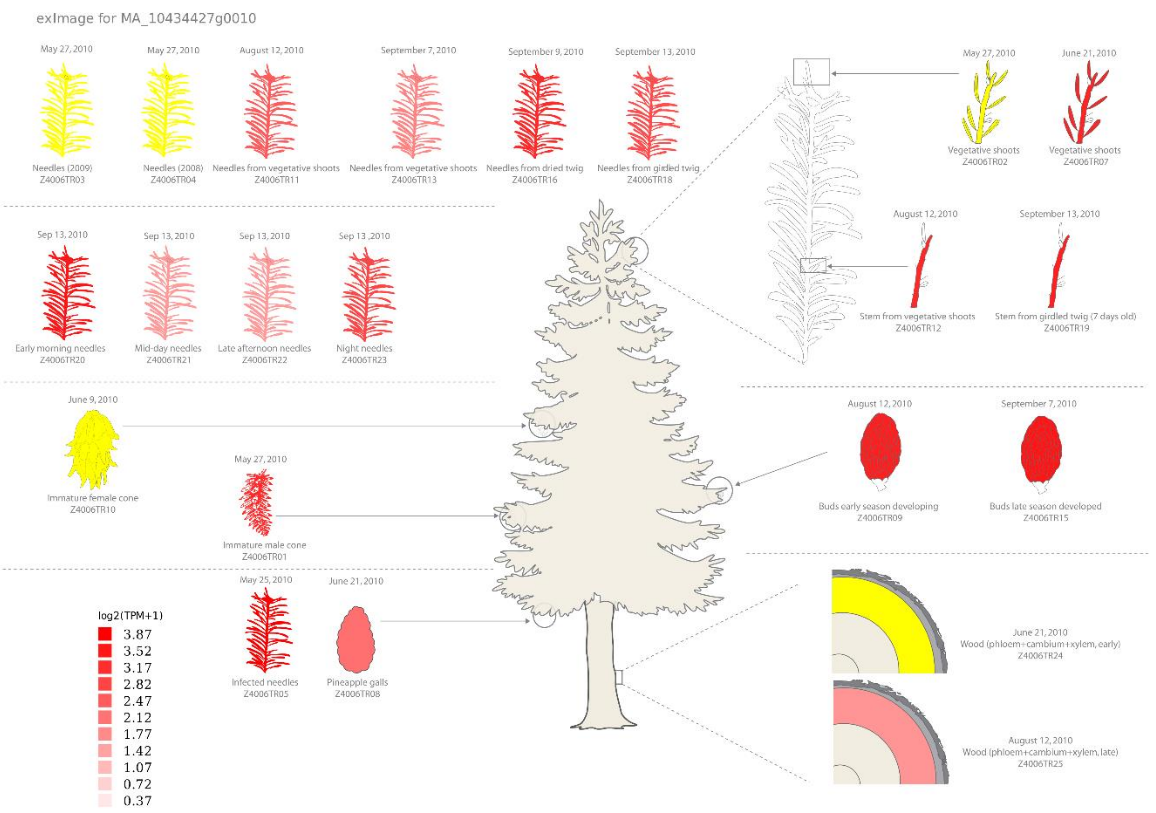


**Fig. S1** The PaAOX1 expression profile in Plantgenie.org database. Expression is highest in the needles and vegetative shoots. Some expression is present also in the wood sample (containing phloem, cambium, and xylem) harvested in August.

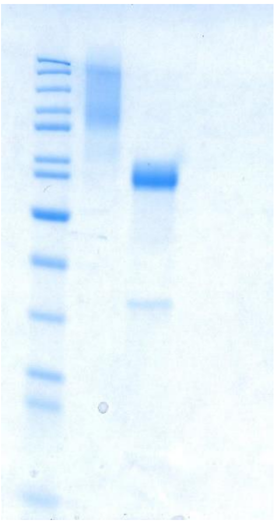

**Fig. S2** The PaAOX1 protein was produced in Pichia pastoris and purified using Ni-NTA affinity chromatography. The purified recombinant PaAOX1 protein shows a high level of glycosylation (lane 2). The purified protein was deglycosylated with PNGaseF (lane 3) and is approximately of the predicted size (61 kDa). The 36 kDa band visible in lane 3 is the PNGaseF enzyme used in deglycosylation.

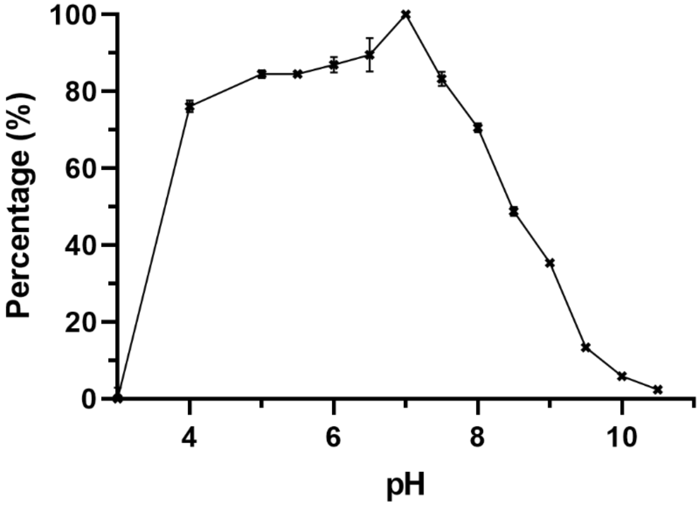


**Fig. S3** Optimal pH for PaAOX1 activity was determined using HRP-coupled ABTS assay using glycerol as a substrate. Activity of PaAOXx1 was compared to the highest activity obtained at pH 7 and shown from pH 3 to 10.5. Activities from pH 3 to 7.5 were measured with McIlvaine buffer, from pH 8 to 9 with Tris buffer, and from pH 9.5 to 10.5 with glycine-NaOH buffer.

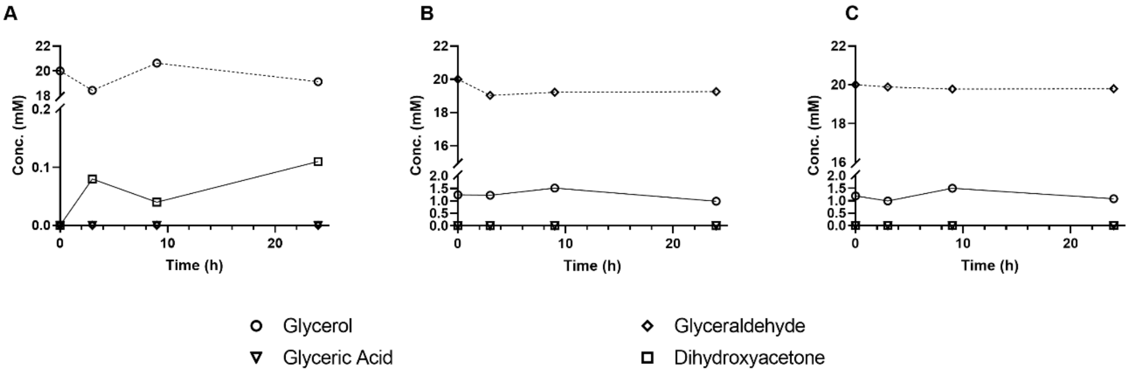


**Fig. S4** Control reactions contained boiled PaAOX1 instead of active PaAOX1. The reactions were followed for 24 h. Substrates: A) Glycerol B) D-glyceraldehyde C) L-glyceraldehyde. No replicates were made for the control reactions.

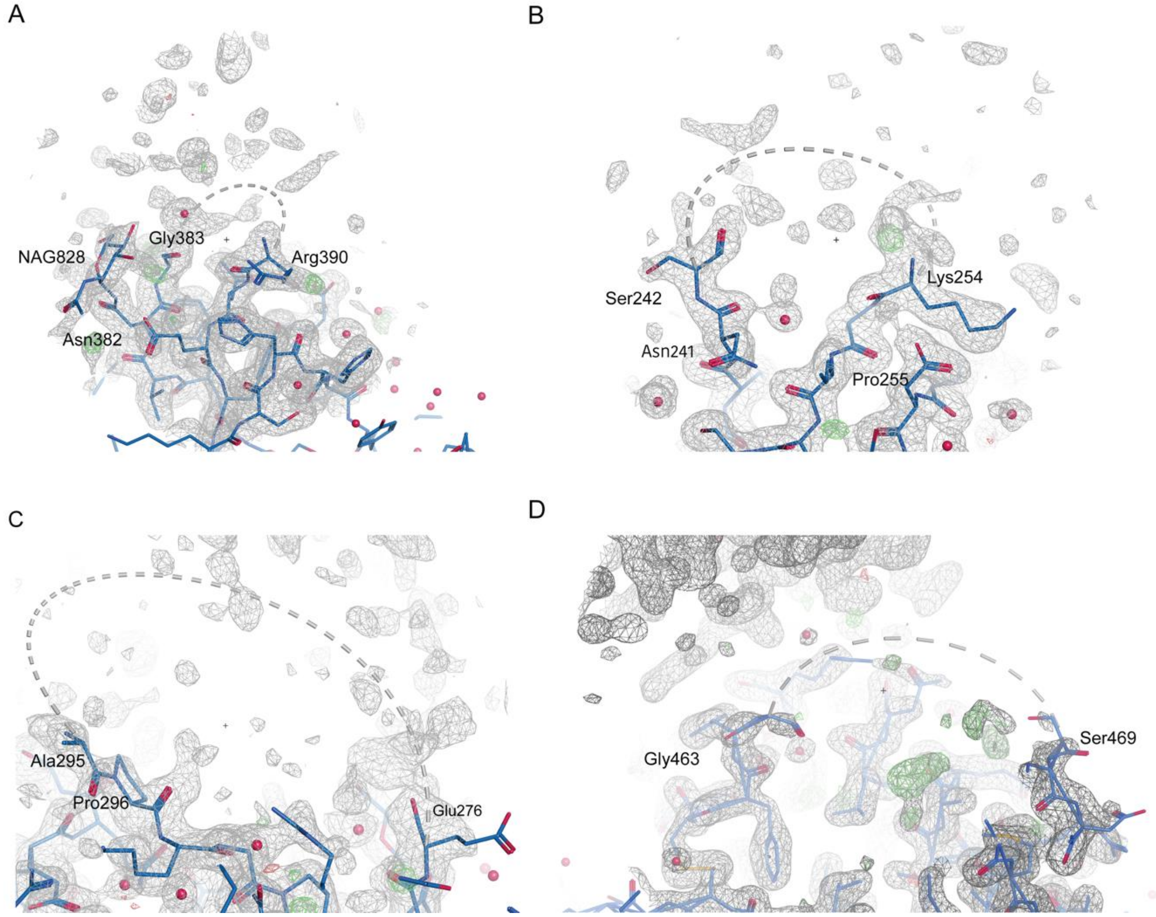


**Fig. S5** Loops of PaAOX1 with missing density. Each of the non-crystallographic symmetry (NCS) PaAOX1 molecules (blue) contain at the surface of the protein three loops, which have a missing density. The loops of the NCS molecule A are shown in picture A to C, D shows a loop which only lacks density in the B molecule. The last amino acids are shown as sticks and overlaid with the final electron density map (gray mesh, sigma level of 1 RMSD). A) The loop closest to the active site with a NAG (N-acetyl D-glucosamine) attached to the Asn382. The loops shown in b) and c) are part of the FAD (flavin adenine dinucleotide) binding domain. D) shows a loop close to the C-terminus which lacks density only in molecule B.

**Table S1** Compounds tested as substrates for PaAOX1 with no detectable or trace activity. The activity assay was conducted at 30 °C for 1 hour with 10 mM substrate in Na-phosphate buffer, pH7.

| **Mono-, di- and trisacchharides** | **Activity** |
|---|---|
| Arabinose | N.D. |
| Cellobiose | N.D. |
| Galactose | trace |
| Glucose | N.D. |
| Lactose | N.D. |
| Maltose | N.D. |
| Mannose | N.D. |
| Raffinose | N.D. |
| Rhamnose | N.D. |
| Xylose | N.D. |
| **Primary alcohols** | |
| Methanol | N.D. |
| Ethanol | N.D. |
| Butanol | trace |
| Pentanol | N.D. |
| Hexanol | N.D. |
| Octanol | N.D. |
| Nonanol | N.D. |
| Decanol | N.D. |
| Dodecanol | N.D. (not fully soluble) |
| Hexadecanol | N.D. (not fully soluble) |
| **Secondary and aromatic alcohols** | |
| 2-propanol | N.D. |
| Benzyl alcohol | N.D. |
| 4-methoxybenzyl alcohol | N.D. |
| **Polyols** | |
| Arabitol | N.D. |
| Lactitol | N.D. |
| Mannitol | N.D. |
| *Myo*-inositol | N.D. |
| Sorbitol | N.D. |
| Xylitol | N.D. |
| **Acids** | |
| Glycolic acid | N.D. |
| 16-Hydroxyhexadecanoic acid | N.D. (not fully soluble) |
| **Complex carbohydrates** | |
| Arabinogalactan | N.D. |
| Arabinoxylan | N.D. |
| Galactoglucomannan | N.D. |
| Galactomannan | N.D. |
| Galacturonan oligosaccharides Mixture DP10/DP15 | N.D. |
| Glucomannan | N.D. |
| Pectin (citrus peel) | N.D. |
| Xylan | N.D. |
| Xyloglucan | N.D. |
| Xyloglucan oligos | N.D. |
| **Others** | |
| Glycerol-3-phosphate | N.D. |
| Glyceraldehyde-3-phosphate | N.D. |
| 5-hydroxymethylfurfural | N.D. |
| Ethylene glycol | N.D. |

**Table S2** Primers used in this study

| Name | Sequence 5'-->3' |
|---|---|
| PaAOX1_UTR_F | CGAAATCGAACCACATCTTC |
| PaAOX1_UTR_R | CGAATATTGCTTCTACGAATGC |
| PaAOX1pp_F | CTTAGGTACCAATCAGCTCTTGTTCTCTTCTCAG |
| PaAOX1pp_R | CTTATCTAGACATTCAGCCTGCTCTTGAAGAA |

**Table S3** Diffraction data and refinement statistics for the PaAOX1 structure (PDB code 8S2Y)

| Data collection and refinement | |
|---|---|
| Data collection | |
| Space group | P 1 21 1 |
| Unit-cell parameters | |
| Å | a=63.1 b=99.3 c=87.96 |
| ° | α= 90 β=100.3 γ=90 |
| X-ray source | Diamond, I03 |
| Wavelenght (Å) | 0.9537 Å |
| Resolution (Å) | 1.64 |
| Resolution range (Å) | 65.24 - 1.64 |
| Completeness | 94.6 |
| Total No. of observations | 770876 |
| Unique reflections | 123313 |
| ⟨I/σ(I)⟩ | 10.5 |
| Rmerge | 0.1188 (1.5297) |
| Multiplicity | 5.9 |
| Structure refinement | |
| Rwork/Rfree (%) | 17.7/23.7 |
| R.m.s.d., bond distances (Å) | 0.0118 |
| R.m.s.d., bond angles (°) | 1.968 |
| No. of amino-acid residues | 7853 |
| No. of water molecules | 662 |
| No. of ligands | 380 |
| No. of ions | 3 |
| Ramachandran plot‡ | |
| Most favored regions (%) | 95.20 % |
| Allowed regions (%) | 3.86% |
| Disallowed regions (%) | 0.94 % |
